# Spatial structure may favor or disfavor microbial coexistence

**DOI:** 10.1101/2022.08.01.502385

**Authors:** Alexander Lobanov, Samantha Dyckman, Helen Kurkjian, Babak Momeni

## Abstract

Microbes often exist in spatially structured environments and many of their interactions are mediated through diffusible metabolites. How does such a context affect microbial coexistence? To address this question, we used a model in which the spatial distributions of species and diffusible interaction mediators are explicitly included. We simulated the enrichment process, examining how microbial species spatially reorganize and how eventually a subset of them coexist. In our model we find that slower motility of cells promotes coexistence by allowing species to co-localize with their facilitators and avoid their inhibitors. We additionally find that a spatially structured environment is more influential when species mostly facilitate each other, rather than when they are mostly competing. More coexistence is observed when species produce many mediators and consume some (not many or few) mediators, and when overall consumption and production rates are balanced. Interestingly, coexistence appears to be disfavored at low diffusion constants of diffusible mediators, because they lead to weaker interaction strengths. Overall, our results offer new insights into how production, consumption, motility, and diffusion intersect to determine microbial coexistence in a spatially structured environment.

## Introduction

A fundamental question in community ecology is how species coexist [1–3]. The observed diversity in many ecosystems has been tantalizing, given the differences among species and competition forces that could exclude the weak in the struggle for existence. Nevertheless, coexistence is widespread in nature, and microbial coexistence is no exception. From anaerobic granules to microbial mats to gut microbiota, microbes are often found not as isolated populations, but in persistent coexistence with others. Explaining what allows such coexistence is of interest, because of the impact of coexistence in important microbial functions across many industrial, environmental, and health-related processes.

Spatial structure and organization have been long recognized as critical factors shaping coexistence [4–10]. Specific mechanisms through which spatial structure may influence coexistence has also been investigated. One common theme in these mechanisms is the role of space in modulating the interactions among individuals [11]. For example, spatial organization may balance out the impact of biotic and abiotic environmental gradients [12,13]. Alternatively, in a spatially structured environment where progeny are more likely to be in the vicinity of parents, intensified intrapopulation competition can give less competitive species a chance to survive [14]. The interplay between dispersal and competition can also allow coexistence between species that are more competitive growers and species that are better at dispersing and colonizing [15].

Although general concepts of coexistence are expected to apply equally to microbes, microbial communities are unique because of the scale and multiplicity of microbial interactions. Many microbial interactions are mediated via diffusible metabolites—including resources and metabolic byproducts. Despite these unique aspects, some general trends in the role of spatial structure on coexistence appear to hold in microbial communities. For example, spatial structure can create environments with higher potential for coexistence between many types of organisms [6]. Likewise amongst bacterial taxa, spatial structure has been observed to stabilize multispecies bacterial communities by balancing the positive and negative interactions [11]. However, it is interesting to note that in the same report [11], spatial structure with certain parameters is detrimental to coexistence, presumably because it weakens some of the interactions necessary to maintain coexistence.

Spatial heterogeneity has been invoked as a mechanism for microbial coexistence in several contexts, from pioneering work by Gause [16] to more recent examples [10,17]. However, prior reports that address coexistence of metabolically interacting microbes in a spatially structured environment are scarce. In an implicit model, Murrell and Law have shown in a modified Lotka-Volterra model that different length scales of inter- and intra-specific competition can lead to species coexistence in a spatially structured environment [18]. A notable example of explicit modeling of space is the recent work by Weiner *et al*, in which coexistence was examined in territorial populations interacting through diffusible mediators [19]. Our model is distinct from their work in that we allow overlap and dispersal of populations through the shared space. Our motivation is to capture situations in which microbes can disperse inside a matrix that defines the spatial structure. An example of this is the mucosal layer of the digestive or respiratory tract, in which stratification is possible, yet the distribution of different species populations can overlap. Another example is in yogurt or cheese, where spatial structure exists, but populations are not territorial. We modify a previously developed mediator-explicit model [20] to account for spatial structure and the dispersion of species in the same space. Here, we limit our study to one-dimensional (1D) spatial structure as a starting point. We examine in our model conditions under which coexistence is favored.

## Results

### A spatial mediator-explicit model of microbial communities

In our mediator-explicit model, species interact through metabolites that they produce and/or consume (Fig S1) [20]. Each species can produce a subset of metabolites and consume a subset of metabolites. Each of the metabolites in the shared environment can in turn influence any of the species by increasing or decreasing their growth rate (i.e. facilitation or inhibition, respectively) compared to how each species grows in the absence of interactions [20]. We also assume that different interaction mediators additively influence the overall growth rate of each species (see “Model description” in Methods).

We assume a 1D spatial structure which preserves the spatial context but allows the diffusion of metabolites and dispersal of species. Multiple metabolites and species can be present in a single location. Both metabolite diffusion and species dispersal are modeled as random walk processes, characterized with a diffusion coefficient and a dispersal coefficient, respectively. In a typical simulation, we start from an initial distribution in which populations occupy adjacent, non-overlapping spatial locations at low initial density. This choice is made to impose a reproducible initial condition that emphasizes the role of space. Each simulation starts with a network of interactions in which interaction strengths, production and consumption links, and production and consumption rates are assigned randomly. The initial pool typically contains 10 species and 5 interaction mediators. We simulate community enrichment through rounds of growth and dilution [20,21] for 100 generations, and assess the richness of each resulting community (i.e., the number of species stably persisting in the community). We have chosen 100 generations of growth, because we have observed that often this is enough to reliably decide which species stably persist in the community. At each dilution step, we assume that the overall spatial distribution of the community is preserved and all populations at all locations are diluted with the same factor. We recognize that this assumption is not universally true; however we adopt it as the least biased possibility, in the absence of additional information about a particular community. We use a well-mixed version [20]—devoid of any spatial context—with a similar set of parameters for all comparisons. Fig S2 shows an example of the population distributions and dynamics during the course of enrichment. In a simple example, we show that interactions and subsequently the population dynamics are affected by growing in a well-mixed versus spatial environment (Fig S3). We explored the impact of the overall spatial extent of the community and found that within an order of magnitude of change, the outcomes remained the same (Fig S4).

The shift from interspecies competition to intraspecies competition can favors coexistence in a spatially structured environment. To assess this impact, we imposed a cap on total cell number at each location in space. As this cap became more restrictive, it suppressed the most competitive species and led to more coexistence (Fig S5). Since our focus in this manuscript is the impact of interspecies interactions, in the rest of this manuscript we pick the total cell number cap at a level (*k*_*Y*_ = 10^9^ cells/ml) that minimizes the impact of imposed intrapopulation competition.

### A spatial environment favors coexistence more when facilitation among species is prevalent

We first examined how the prevalence of facilitative versus inhibitory interactions impacted coexistence in spatial communities. In our simulations, we dictated the ratio of facilitative and inhibitory interactions in the initial pool of species. Our results show that, similar to a well-mixed environment, more facilitative interactions lead to higher richness in communities that emerge from enrichment (Fig 1, along the x-axis). Additionally, we observe that spatial communities show more coexistence than well-mixed communities when facilitation among species is prevalent (Fig 1, spatial versus well-mixed). Our explanation is that species locally grow better when adjacent to a facilitative partner and grow worse when in the vicinity of an inhibitory partner. The resulting spatial self-organization in effect amplifies facilitative interactions and dampens inhibitory interactions, leading to more coexistence. This is supported by our data which shows that the position of specific species with respect to other species that facilitate or inhibit it can impact the population dynamics (Fig S6). Because of the marked impact of the fac:inh ratio, moving forward, we will examine three conditions, with equal fractions of facilitative and inhibitory influences (fac:inh = 50:50), mostly inhibitory (fac:inh = 10:90), or mostly facilitative (fac:inh = 90:10) to scope the impact on coexistence.

**Figure 1.**
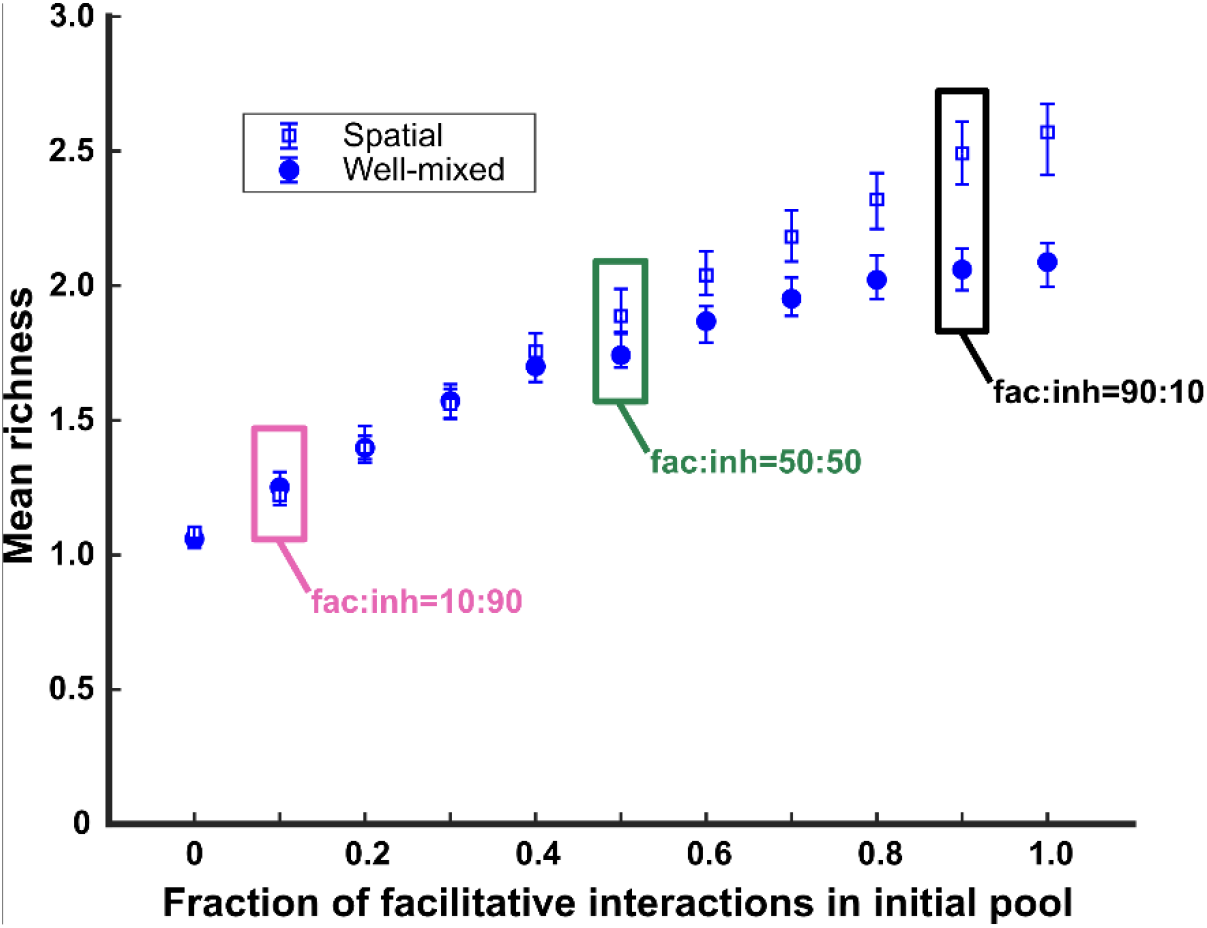
A spatial environment favors coexistence more when facilitation among species is prevalent in the initial species pool. Simulations were run at different fac:inh ratios for both spatial communities (green square) and well-mixed communities (pink circles). Each ratio was run 500 times with the richness (number of species stably surviving at the end of a simulation) averaged over all the simulations. Each simulation started with 10 species and 5 mediators and ran for 100 generations. The error bars are 95% confidence intervals generated by bootstrapping 100 samples. Here, the species dispersal coefficient is 5 × **10**^**−9**^ cm^2^/hr. Boxes mark fac:inh ratios used in later simulations.

### Coexistence is favored when many metabolites are produced and influence an intermediate number of species

Because metabolites are at the center of interspecies interactions in our model, we examined their impact on spatial coexistence of the average number of metabolites produced by each species and the average number of species influenced by each mediator. We found that coexistence is favored when the number of metabolites produced is larger (Fig 2, along the y-axis). This effect was stronger when the metabolite influences were mostly facilitative (fac:inh = 90:10, versus 50:50 or 10:90). In contrast, coexistence achieved its maximum values at intermediate ranges of mediator influence (Fig 2, x-axis), i.e. lower coexistence was observed when each mediator influenced too many or too few species on average. We note that these trends were largely the same between spatial and well-mixed communities.

**Figure 2.**
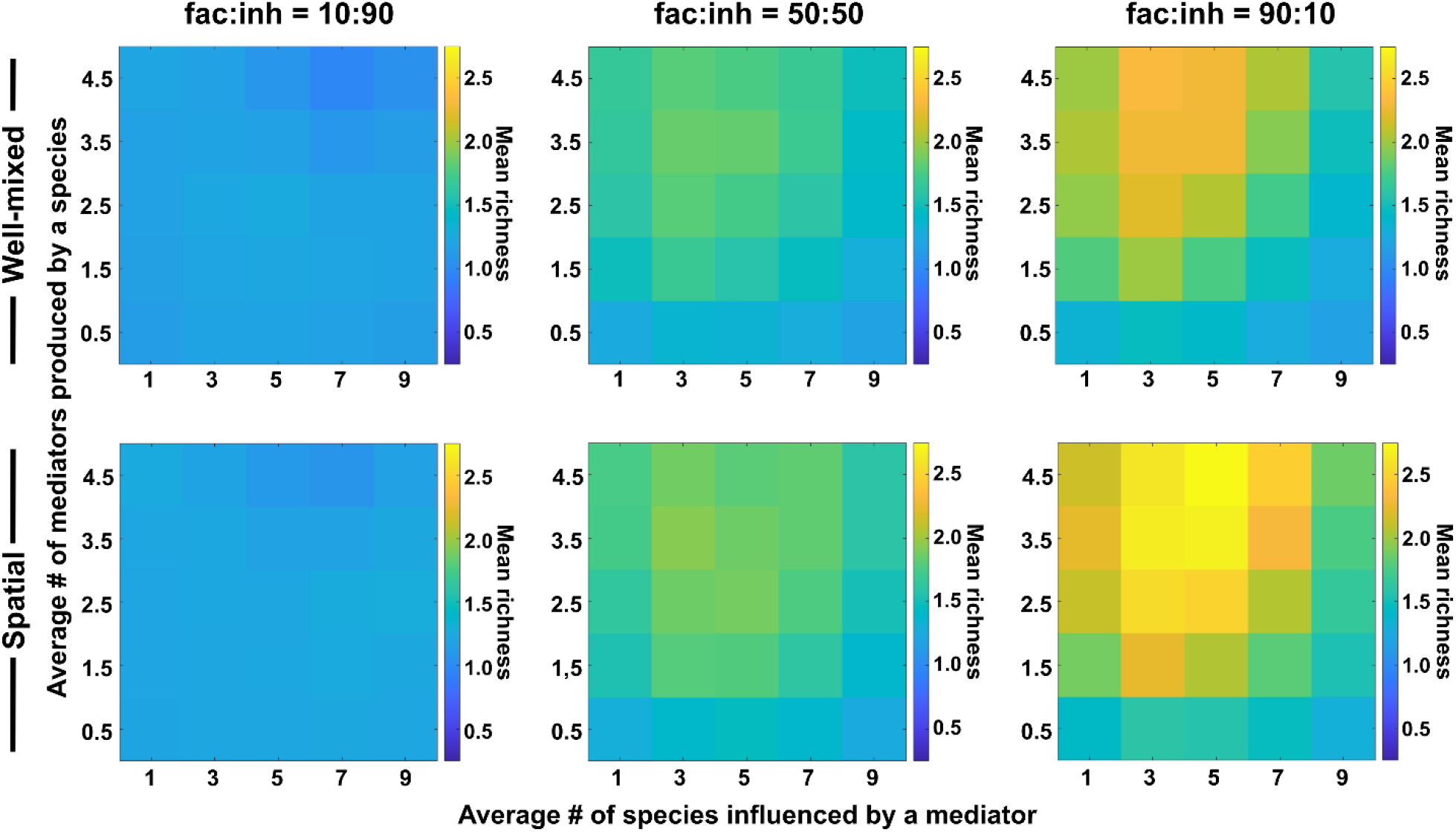
The number of metabolites produced and number of species influenced affect coexistence in spatial and well-mixed communities. Different ranges of production and mediator influence values were analyzed for both well-mixed and spatial communities at three different fractions of fac:inh influences in the initial pool of species (10:90, 50:50, and 90:10). Mean richness (i.e. average number of species surviving at the end of a simulation) was calculated for 500 simulated instances and marked on the color bar. Each simulation started with 10 species and 5 mediators and ran for 100 generations. The x-axis represents the average number of species influenced by a mediator and the y-axis represents the average number of mediators produced by each species.

Our explanation is that a larger range of production offers more opportunities for interaction, which through the enrichment process lead to the selection of facilitative subsets that coexist [20]. A low mediator influence range works in the opposite direction, reduces opportunities for interactions and results in lower coexistence. Very high mediator influence range potentially leads to more self-facilitation (i.e. producing a metabolite that is beneficial to the producer species), which our data suggests can lead to take-over by a single species and a lower coexistence as a result (Fig S7).

### Coexistence is higher when there is balance between production and consumption of mediators

We next asked how the rates of production and consumption of mediators would influence coexistence. To address this question, we surveyed a range of average rates of production and consumption. We observed that the highest levels of coexistence occurred when there was a balance between consumption and production rates among species, with slightly higher production than consumption (Fig 3).

**Figure 3.**
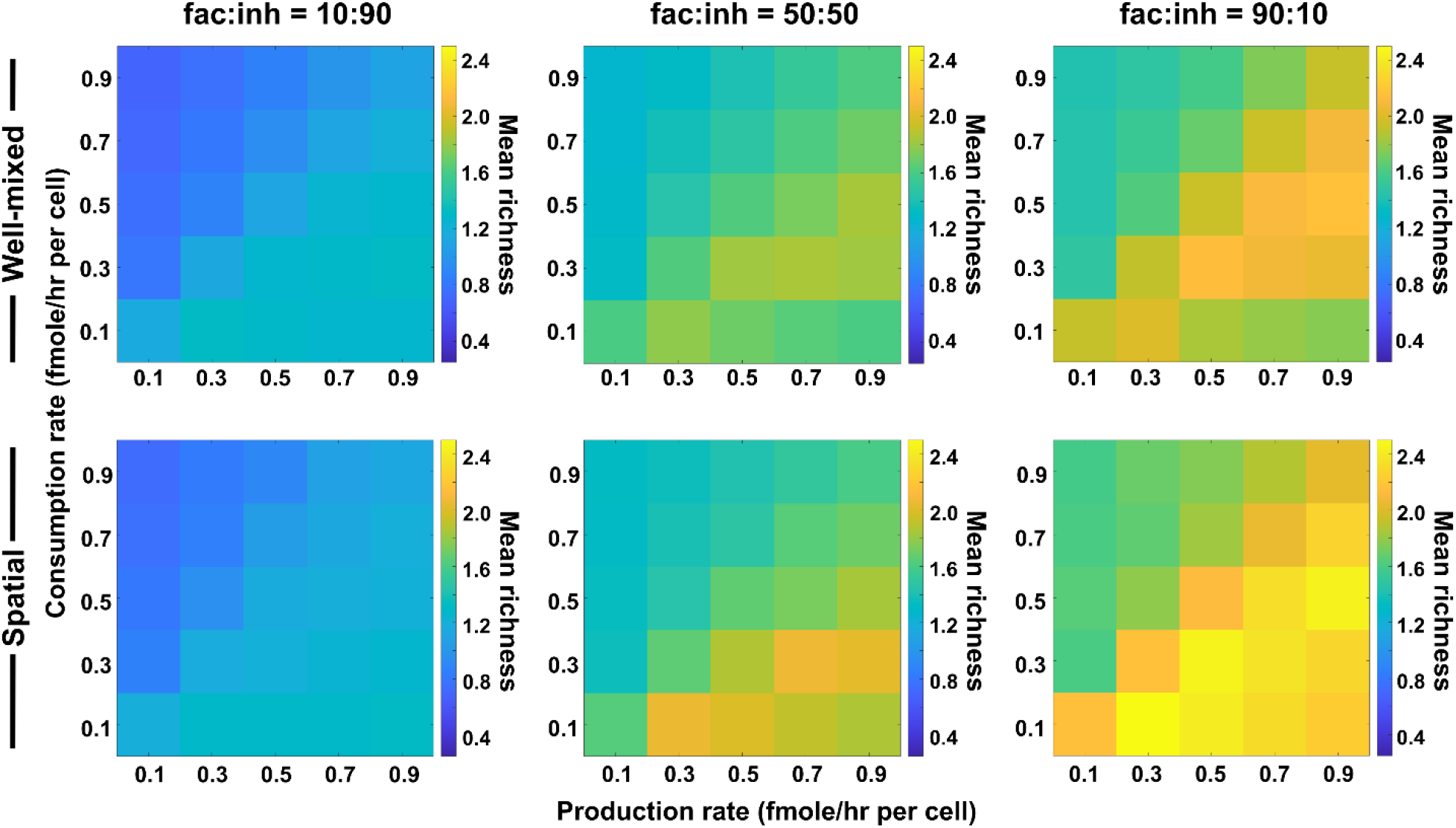
Coexistence is higher where there is a balance between production and consumption of mediators. Different average production and consumption rates were analyzed for both well-mixed and spatial communities at three different fractions of fac:inh influences in the initial pool of species (10:90, 50:50, and 90:10). Mean richness (i.e. average number of species surviving at the end of a simulation) is calculated for 500 simulated instances. Each simulation started with 10 species and 5 mediators and ran for 100 generations. Color bar represents the average richness. The x-axis represents the average production rate of mediators and the y-axis represents the average consumption rate of mediators.

Our justification for the observed pattern is that in one extreme where production is too high (lower right corner of each plot), mediators will build up in the environment. This will put the community in a regime that consumption is not enough to create a feedback, i.e. “reusable mediators” as discussed in [20], which leads to lower coexistence. In the other extreme, when consumption is too high (upper left corner of each plot), metabolites that mediate the interactions will be depleted from the environment, leading to an effectively weaker interaction and thus lower coexistence. However, when production is slightly higher than consumption, metabolite quantities are sufficient to create strong interactions and facilitation feedbacks, leading to higher coexistence. While coexistence is slightly higher in the spatial communities compared to well-mixed ones, the production-consumption trends apply equally to spatial and well-mixed communities, as expected.

### Limited species dispersal in a spatial environment allows more coexistence, especially when facilitation is common

Because species dispersal is a major factor in preserving community spatial structure, we examined how the dispersal coefficient affected coexistence outcomes. For this, we kept the diffusion coefficient of the mediators fixed and surveyed mean richness among many instances of communities randomly assembled (n = 500). When the diffusion coefficient for species approaches zero and cells remain in their original spatial location, we observe higher levels of coexistence (Fig 4). We also observed that the impact of lower dispersal is stronger in communities in which most interactions are facilitative rather than inhibitory.

**Figure 4.**
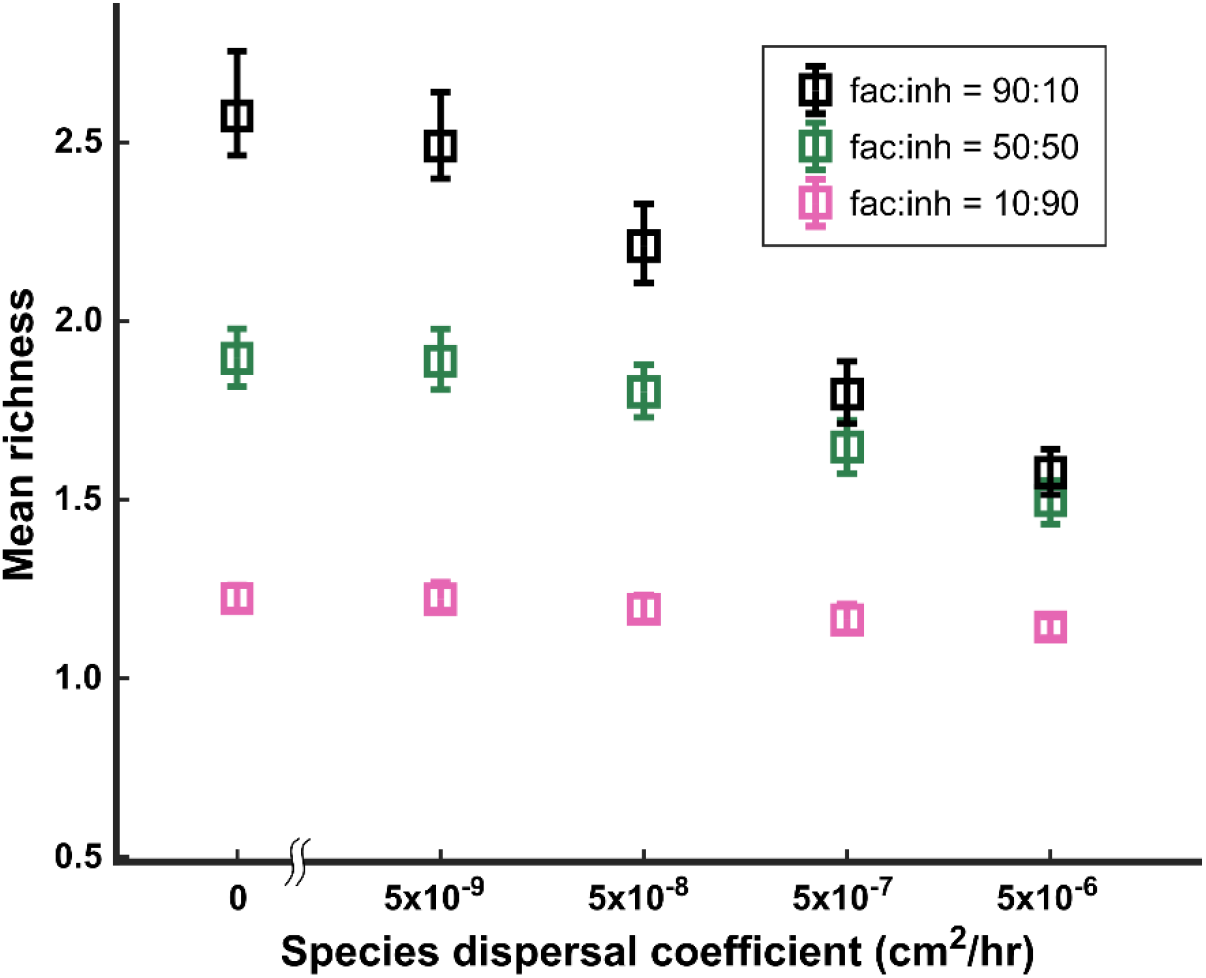
Lower dispersal rates allow more microbial coexistence. Communities in spatially structured environments were simulated with different dispersal coefficients at three different fractions of fac:inh influences (10:90, 50:50, and 90:10). Mean richness (i.e. average number of species surviving at the end of a simulation) was calculated for 500 simulated instances. Each simulation started with 10 species and 5 mediators and ran for 100 generations. The error bars are 95% confidence intervals generated by bootstrapping 100 samples.

Our explanation is that lower dispersal rates mean that species grow best in spatial locations that are more supportive for their growth, which is in the vicinity of their beneficial partners and away from competitors or inhibitors. As discussed in Fig 1, such self-organization effectively amplifies facilitative interactions and de-emphasizes inhibitory interactions, leading to a higher coexistence. This is also consistent with the observation that the effect of dispersal rate is strongest when the proportion of facilitative interactions is highest. As the dispersal coefficient increases, the self-organization gets washed away by dispersal of cells to less than ideal locations for their growth and its benefit for coexistence diminishes.

### Coexistence is disrupted when the diffusion of mediators is too slow

The rate of diffusion of metabolites also has the potential to affect coexistence. We investigated coexistence over a range of mediator diffusion coefficients. We still typically observe a higher mean richness for spatial communities compared with the well-mixed communities (Fig 5). However, unlike the conventional wisdom, as the diffusion of mediators becomes slower, coexistence in spatial communities decreases. At low diffusion coefficients, coexistence drops even below that of corresponding well-mixed communities. We associate this trend to weaker effective interactions among species at lower diffusion coefficients. Mediators that are involved in facilitation play a major part in allowing coexistence of species [20]; if these mediators get consumed by nearby species and do not travel long enough to reach other members of the community, the interaction-driven mechanism of coexistence is disrupted.

**Figure 5.**
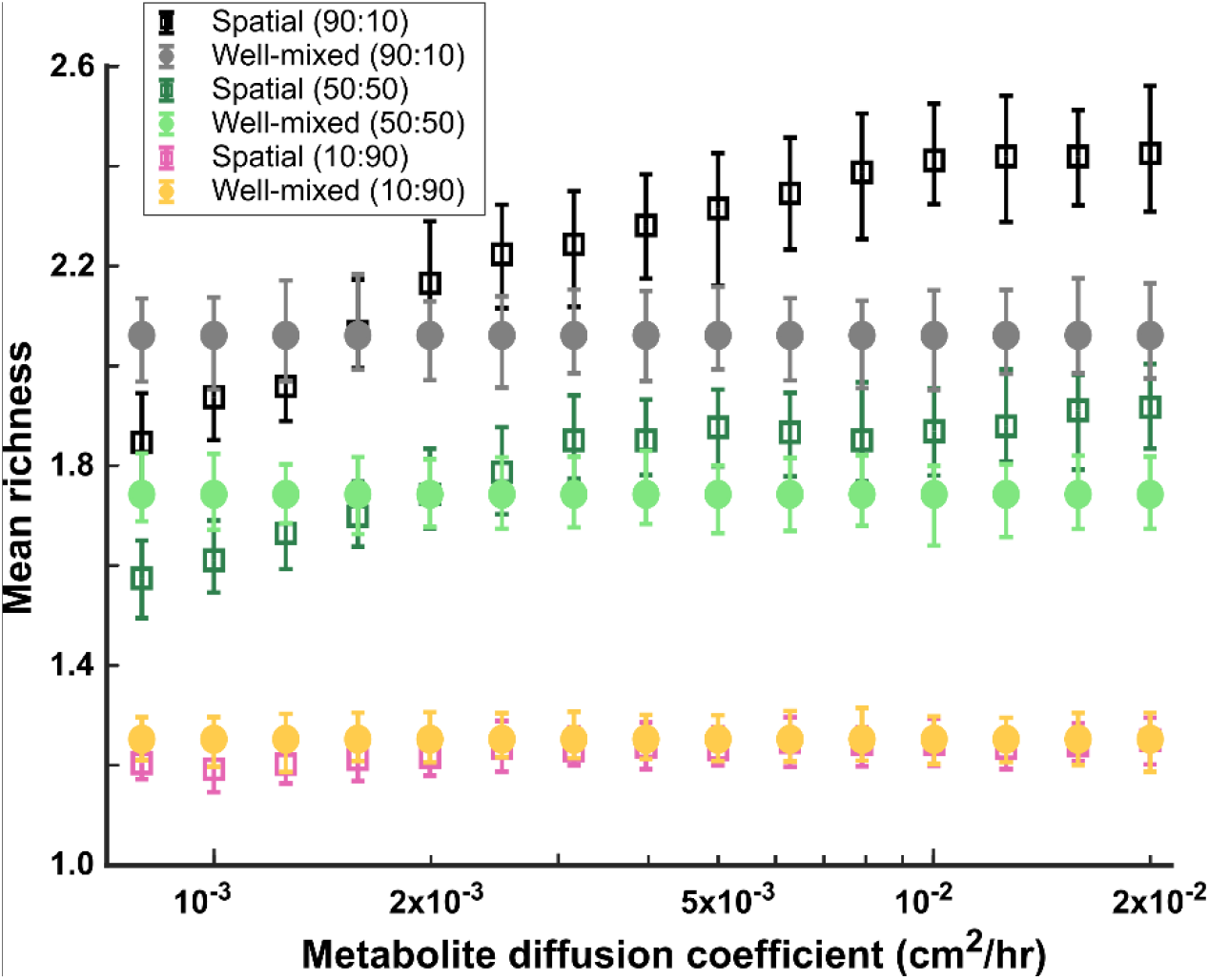
At higher diffusion coefficients of mediators more coexistence is possible. A range of metabolite diffusion coefficients were simulated in spatial communities (squares) at three different fractions of fac:inh influences (10:90, 50:50, and 90:10). We simulated corresponding well-mixed communities (circles) for comparison. Each condition was run 500 times with the richness (number of species stably surviving at the end of a simulation) averaged over all the simulations. Each simulation started with 10 species and 5 mediators and ran for 100 generations. The error bars are 95% confidence intervals generated by bootstrapping 100 samples.

## Discussion

Our results dispel the common presumption that a spatially structured environment will universally lead to more coexistence. We find that, compared to a well-mixed environment, a spatial environment can favor or disfavor coexistence depending on the balance between species dispersal and the diffusion of interaction mediators. Interestingly, a lower species dispersal rate favors coexistence, but this effect can be diminished or even reversed if accompanied by low mediator diffusion rates. Coexistence is favored when species have a broad range of consumption and an intermediate range of production of interaction mediators. Additionally, we predict more coexistence when there is a balance between overall production and consumption rates for mediators.

The spatial structure of microbial communities has been extensively studied for example in simulating the development of biofilms [22–25], for specific interactions among species [26–28], or for modeling game-theory dynamics [29–32]. However, the focus on detailed mechanisms make such modeling strategies not suitable for general investigation of coexistence. As expected, the generality of conclusions is partially lost when detailed mechanisms are incorporated [33,34].

We have made assumptions in our model to simplify the configuration and make the analyses and interpretations easier. We asked if making these assumptions more realistic would affect our conclusions. For example, we have assumed no carrying capacity limit for the growth of our populations. We tested if imposing a total population limit, enforced at each spatial location, would alter our conclusions. We saw an impact when a local carrying capacity was incorporated in our model (Fig S4), but chose parameters to minimize this impact. We also tested the impact of the spatial extent of the community (*Z*), and observed that our results were largely unaffected if the community’s spatial extent was changed by an order of magnitude (Fig S5). The effect of larger changes in the spatial extent can be examined by scaling the diffusion and dispersal coefficients accordingly.

Beyond the details of our assumptions, there are also alternative representations of interactions among species, including a simplified Lotka-Volterra model and its variations [35], a consumer-resource model [21,36], or a reduced metabolic model [37]. There are trade-offs in tractability and complexity in choosing which model to use. Our reasoning for adopting the mediator-explicit model was to (1) explicitly include metabolites that mediate the interactions in the model [38], and (2) incorporate both metabolites that support the growth of other species as well as those that are inhibitory, such as waste products and toxins [38]; and (3) keep the model simple to allow a clear interpretation of mechanisms and processes when analyzing the results. We think it will be worthwhile to compare the predictions of other models to clarify what assumptions are necessary to generate the trends we have obtained and how general the conclusions are.

If spatial organization of cells matters, we also expect that the initial spatial position of species in the community impacts coexistence. To test this, we started from 100 simulations instances and in each case, we tried 100 rearrangements, each obtained by shuffling the spatial position of species, while keeping the species properties and interactions intact. Interestingly, in many cases coexistence was affected (Fig S8), indicating that the adjacency to partners is an important determinant of spatial coexistence (as also suggested by Fig S3 and Fig S6). When we examined the effect of the fac:inh ratio on these outcomes, we observed that larger changes in richness when facilitation interactions were more prevalent in the community (Fig S9), which aligns with many of our other results showing that facilitation amplifies the positive effect of spatial structure on coexistence.

Finally, our model assumes that dispersal and diffusion rates are uniform across species and metabolites, respectively. However, dispersal ability can vary widely across microbial taxa, depending on cell size, motility type, chemotaxis, quorum sensing, and other factors. And how the dispersal rates of individuals scale up to affect population and community level dynamics is not well understood. Likewise the diffusion rates of metabolites has the potential to vary greatly with molecule size and shape. Although outside the scope of this work, we are exploring heterogeneity in these rates of movement as an interesting follow-up.

Overall, we believe this work revisits how spatial structure—and spatial self-organization— affects community assembly and coexistence. In our model that emphasizes the contributions of interspecies interactions, we find that the impact of spatial structure on coexistence largely arises from two processes: (1) spatial self-organization, which can improve coexistence by favoring facilitation over inhibition, and (2) localization of interactions, which can promote coexistence in association with self-organization or hamper coexistence by slowing down and weakening species interactions.

## Methods

### Model description

Our model is an extension of a model introduced earlier [20] in which a set of species interact with each other through diffusible mediators. Each mediator is produced by a subset of species, consumed by a subset of species, and has a positive or negative influence on the growth rate of some species (Fig S1).

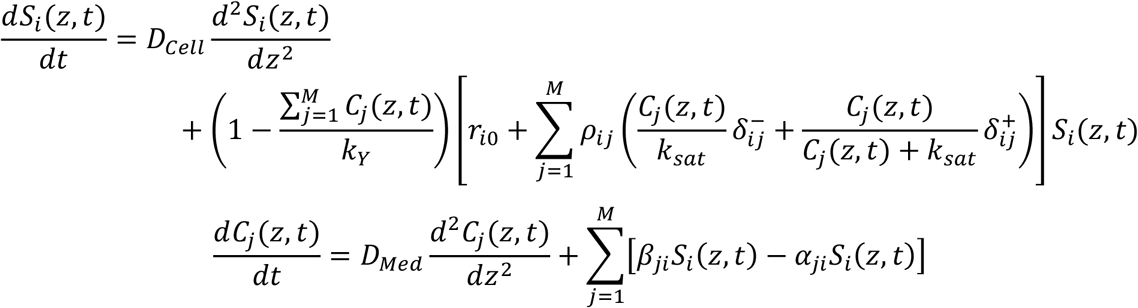

Here *S*_*i*_ is the spatiotemporal distribution of species *i* (*i* = 1, …, *N*_*c*_). *C*_*j*_ is the spatiotemporal distribution of mediator *j* (*j* = 1, …, *N*_*m*_). *D*_*Cell*_ is the dispersal coefficient for cells. *D*_*Med*_ is the diffusion coefficient for mediators. *k*_*Y*_ is the total cell carrying capacity at each location. *ρ*_*ij*_ is the interaction coefficient expressed as the impact of mediator *j* on species *i*. Additionally,

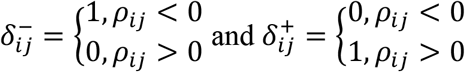

*k*_*sat*_ is the interaction strength saturation level. *β*_*ji*_ and *α*_*ji*_ are average production rate and consumption rates, respectively, between species *i* and mediator *j*.

Typical parameters used in our simulations (unless otherwise stated) are listed in Table 1.

**Table 1.** Parameters used in our simulations are listed.

| Parameter | Value (unit) |
| --- | --- |
| Number of instances examined ($N_s$) | 500 |
| Number of cell types in the initial pool ($N_c$) | 10 |
| Number of interaction mediators ($N_m$) | 5 |
| Total initial cell density ($TID$) | $10^4$ (cells/ml) |
| Interaction strength saturation level ($k_{sat}$) | $10^4$ (cells/ml) |
| Population extinction threshold ($ExtTh$) | 0.1 (cells/ml) |
| Population dilution threshold ($DilTh$) | $10^7$ (cells/ml) |
| Average consumption rate ($\alpha$) | 0.15 (fmole per cell per hour) |
| Average production rate ($\beta$) | 0.1 (fmole per cell per hour) |
| Probability of production link per population ($q_p$) | 0.5 |
| Probability of influence link per mediator ($q_c$) | 0.5 |
| Maximum interaction strength magnitude ($r_{int,0}$) | 0.2 (1/hr) |
| Number of generations for enrichment ($nGen$) | 100 |
| Dispersal coefficient for cells ($D_{Cell}$) | $5 \times 10^{-9}$ (cm<sup>2</sup>/hr) |
| Diffusion coefficient for mediators ($D_{Med}$) | $1.8 \times 10^{-3}$ (cm<sup>2</sup>/hr) |
| Total carrying capacity per dz ($k_Y$) | $10^9$ cells/cm |
| Total community spatial extent ($Z$) | 0.5 cm |
| Spatial resolution for species distributions ($dz$) | 0.005 cm |
| Cell growth update and uptake timescale ($d\tau$) | 0.01 hr |
| Mediator diffusion time-step ($dt$) | $0.1 dz^2 / D_{Med}$ |
| Cell dispersal simulation time-step ($dc$) | $0.1 dz^2 / D_{Cell}$ |

### Model implementation

We solve the equations in ‘Model description’ numerically in Matlab® using a finite difference discrete version of the equations. Mediator diffusion and cell dispersal take place often at very different time scales. To simulate these processes, we use different numerical time-steps to update the mediator and cell distributions. To allow flexibility in modeling different diffusion and dispersal coefficients, we used asynchronous updates with two independent time steps: one for updating the diffusion of metabolites and another one for growth and dispersal of cells. The source codes are shared for transparency and reproducibility (see ‘Code availability’).

To assess coexistence, we use a criterion similar to [20]. In short, any species whose density drops below a pre-specified extinction threshold (*ExtTh*) is considered extinct. Among species that persist throughout the simulation, only those with relative frequencies equal to or larger than 90% of the fastest growing species in the last 20 generations of the simulation are considered to coexist. Species with a declining relative frequency are assumed to go extinct later and are not considered to be part of coexisting communities.

### Statistics

Mean richness values are calculated by averaging the richness values calculated over all simulated instances for a given condition. Confidence intervals for mean richness values are calculated by bootstrapping over all simulated instances for a given condition. The standard routine in Matlab, `bootci`, is used in all cases for bootstrapping.

## Code availability

All the codes used for this manuscript are available at: https://github.com/bmomeni/spatial-coexistence.

## Author contribution

SD, HK, and BM conceived the idea. SD, AL, and BM developed the code. SD, AL, and BM ran simulations and analyzed the data. AL, SD, and BM wrote the manuscript. AL, SD, HK, and BM edited the manuscript.

## Acknowledgements

BM would like to acknowledge the Award for Excellence from Smith Family Foundation that supported this work. BM would like to also thank the biology department at Boston College for their continued support. We would like to acknowledge the support for AL through an Undergraduate Research Fellowship from Boston College.

## Supplementary information

**Fig S1.**
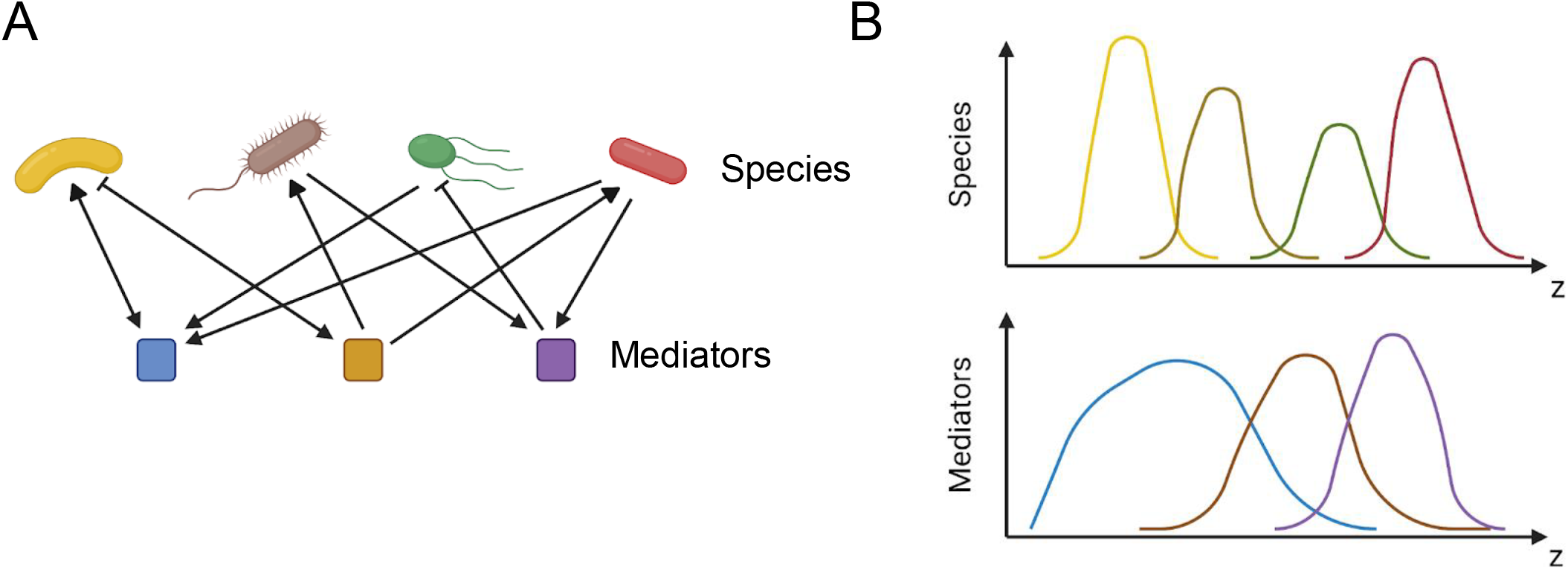
Schematic illustration of the overall setup. (A) Species are engaged in interactions with other species through metabolite that mediate those interactions. Each species produces a subset of mediators and consumes a subset. Each mediator can positively or negatively modulate the growth rate of the species it influences. (B) In a one dimensional (1D) spatial context, species and mediators are defined as functions of space that change over time because of population growth and dispersal as well as mediator production, consumption, and diffusion.

**Fig S2.**
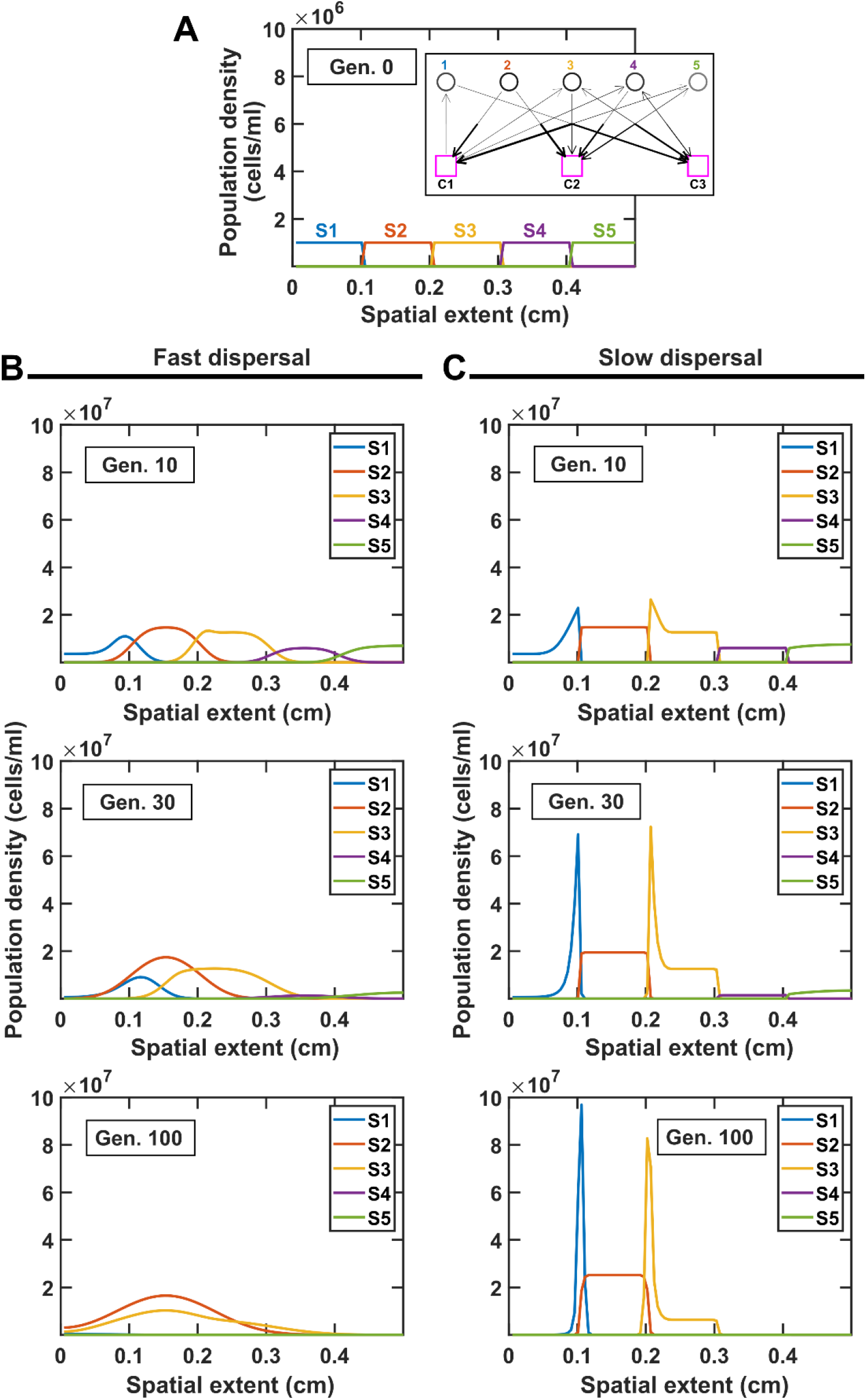
Comparing the spatial distribution of species at different dispersal rates illustrates the impact of dispersal on coexistence. (A) Species are initially stacked over the spatial extent of the community. Here we examine a simple exam with 5 species and 3 mediators (interaction network shown in the inset). Different progressions are observed at high (B, *D*_*Cell*_ = 5×10^™6^ cm^2^/hr) versus low (C, *D*_*Cell*_ = 5×10^™9^ cm^2^/hr) dispersal rates, leading to different coexistence outcome. In both cases, *D*_*Med*_ = 1.8×10^™2^ cm^2^/hr

**Fig S3.**
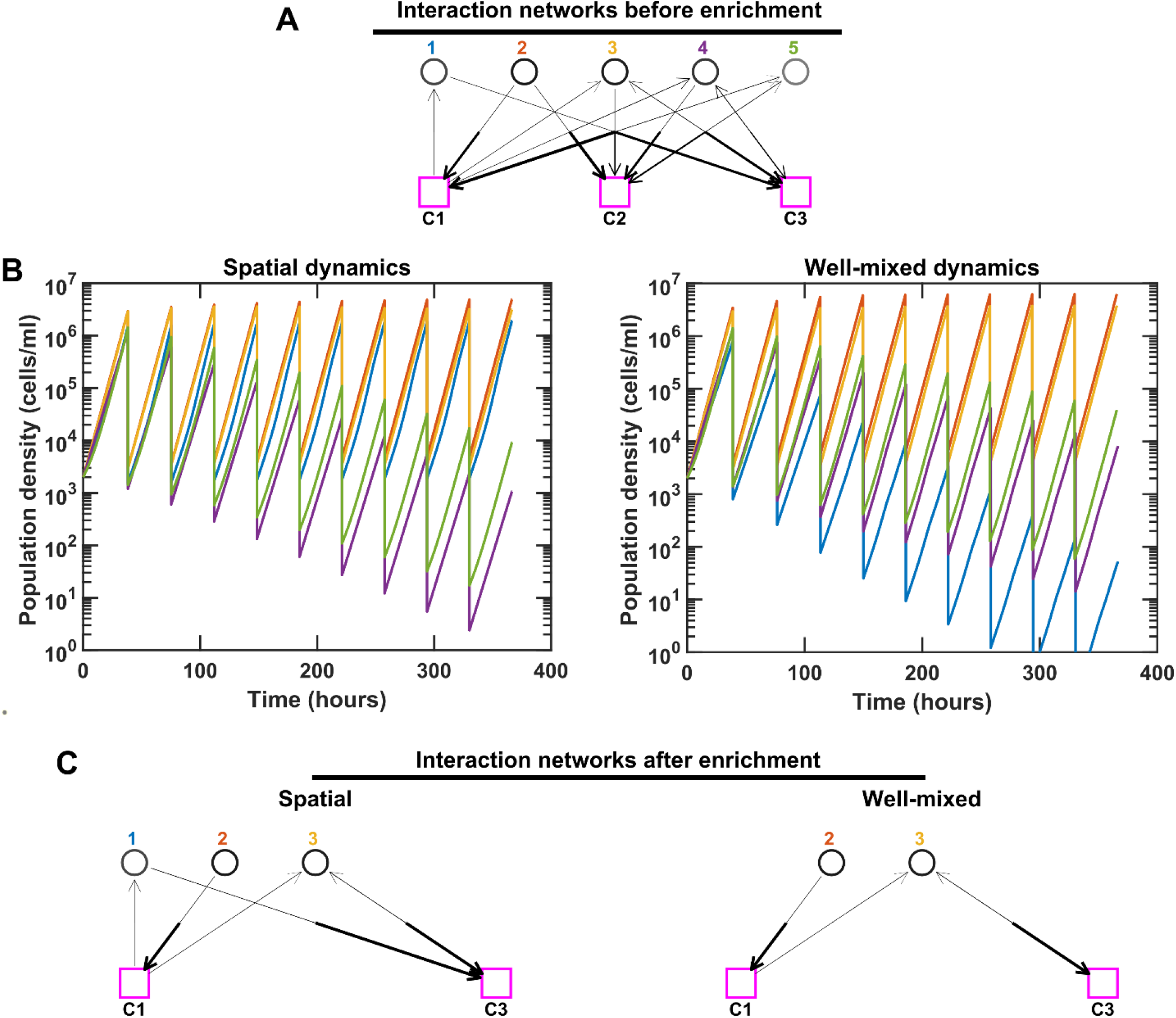
Species interactions and dynamics are different in spatial versus well-mixed environments, leading to different coexistence outcomes. For simplicity, here we consider an initial pool of 5 species with 3 mediators. We keep the same interaction network (A) in both spatial and well-mixed conditions and simulated the species dynamics. For spatial communities, the population density represents the sum of all densities across different spatial extents. We ran the simulation in both cases through 10 rounds of growth and dilution. Different population dynamics and ultimately different coexistence outcomes are observed in these cases after simulating 100 generations. Note that even though species 4 and 5 in the spatial community and species 1, 4, and 5 in the well-mixed community are still present in the final communities, because they exhibit a trend of decline in relative frequency, we consider them not to coexist (in accordance with our coexistence criterion, see Methods). (B-C) In this particular example, proximity of Species 1 to a beneficial Species 2 in the spatial community allows a strong boost in the growth of Species 1, leading to its coexistence with Species 2 and 3. In contrast, Species 1 in the well-mixed community receives a smaller portion of C_1_, not enough to allow Species 1 to keep up with the other species. In C, only coexisting species and mediators related to them are shown.

**Fig S4.**
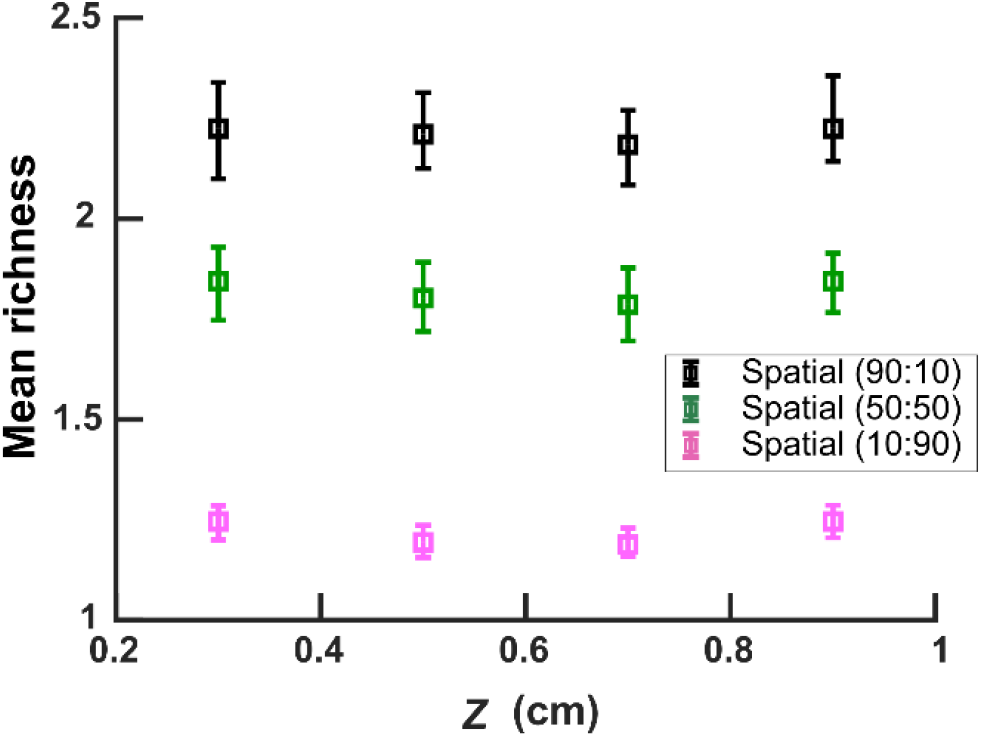
Within the same order of magnitude, community’s spatial extent does not have a large impact on spatial coexistence. All parameters other than community’s spatial extent are kept fixed. Here *D*_*Cell*_ = 5×10^™8^ cm^2^/hr. Legend shows the values of fac:inh.

**Fig S5.**
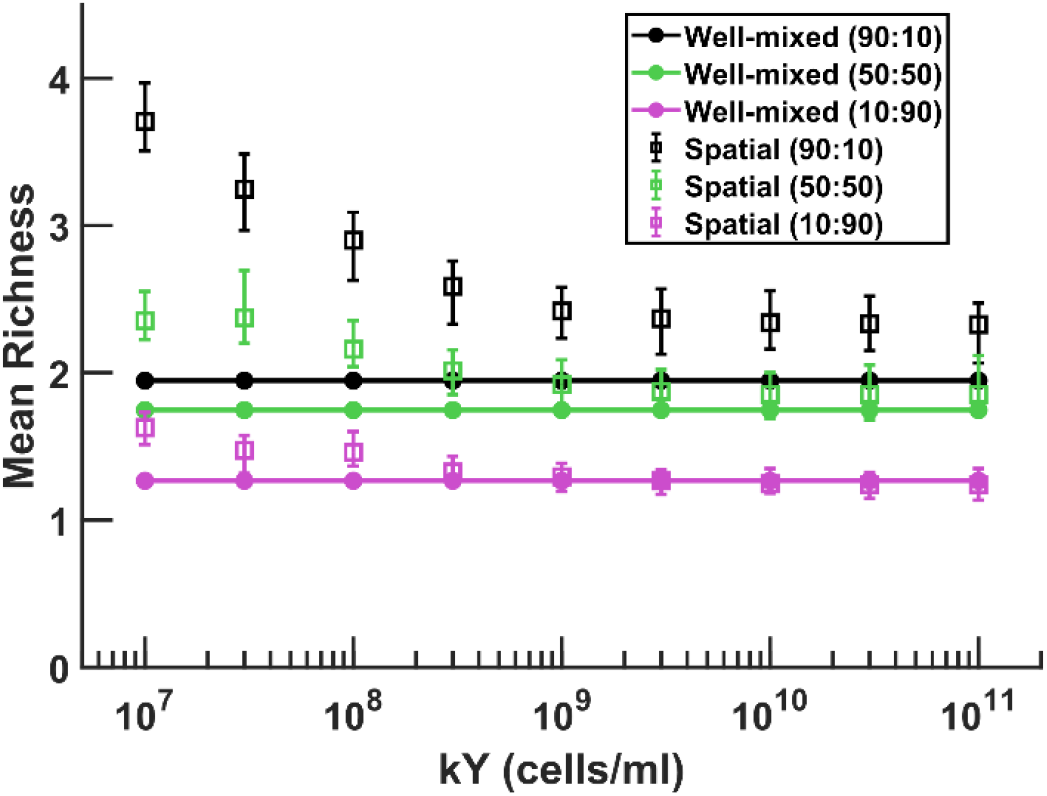
Imposing a local carrying capacity favors species coexistence. All parameters other than the local total carrying capacity kY are kept fixed. Imposing a local carrying capacity on the total cell number (see the equations described in Methods) lowers the competitive advantage of the most successful species and allows more coexistence. To focus on the effect of interspecies interaction, we assume kY = 10^9^ cells/ml in our other simulations to minimize the impact of kY (i.e. a forced intrapopulation competition) on our results.

**Fig S6.**
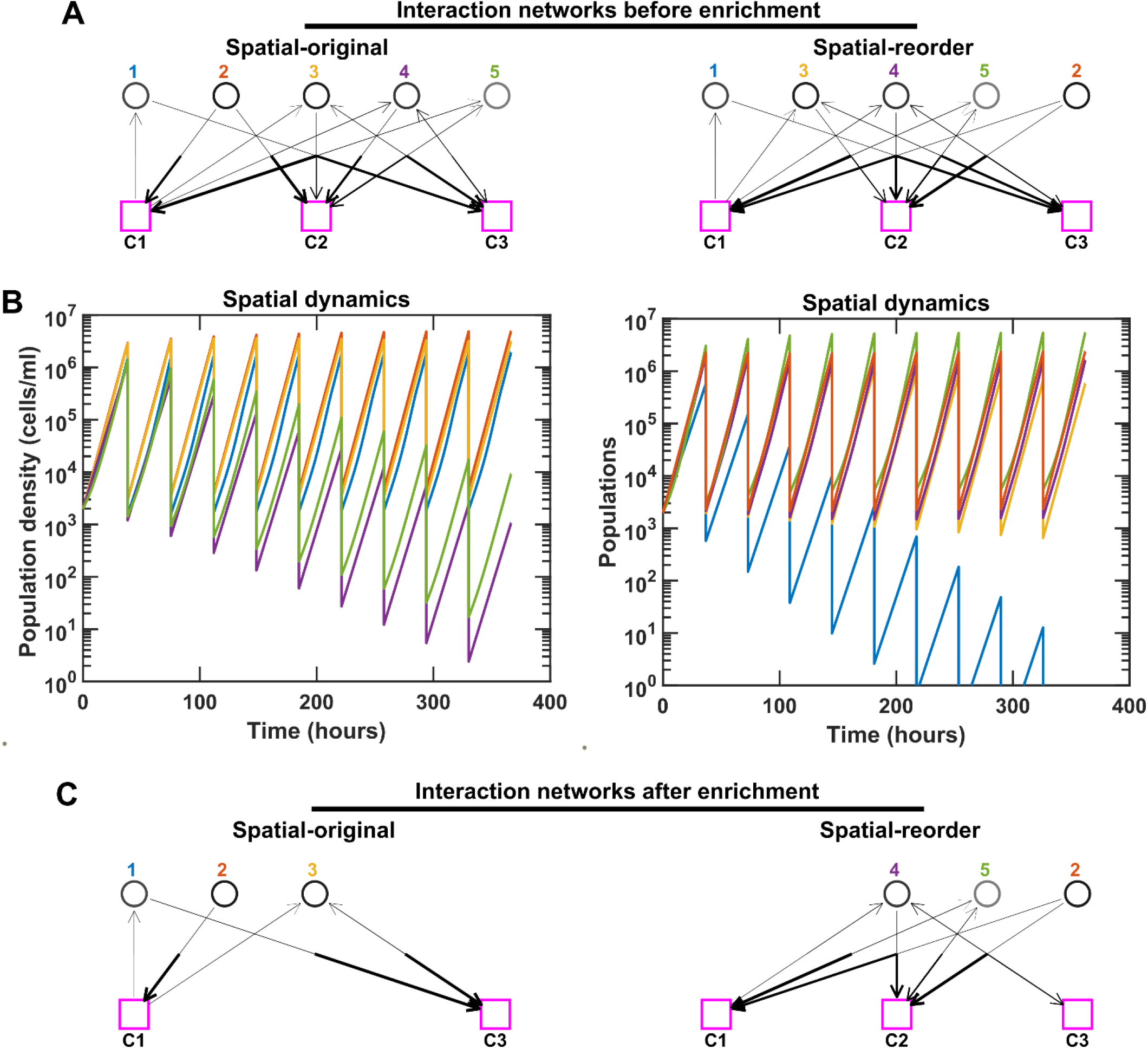
Rearranging the order of species can modulate the strength of interspecies interactions and impact spatial coexistence outcomes. For simplicity, here we use the same examples as Fig S3 with 5 species and 3 mediators in the initial pool. We keep the same interaction network (A), but rearrange the initial order of species in the spatial community (original: 1, 2, 3, 4, 5; reorder: 1, 3, 4, 5, 2). We ran the simulation in both cases through 10 rounds of growth and dilution. Different population dynamics and ultimately different coexistence outcomes are observed in these cases after simulating 100 generations. Note that even though species 4 and 5 in the original community and species 1 and 3 in the reorder community are still present in the final communities, because they exhibit a trend of decline in relative frequency, we consider them not to coexist (in accordance with our coexistence criterion, see Methods). (B-C) In this particular example, after changing the order, Species 1 and 3 receive less of the benefits from Species 2, leading to the emergence of Species 2, 4, and 5 as an alternative stable coexistence. In C, only coexisting species and mediators related to them are shown.

**Fig S7.**
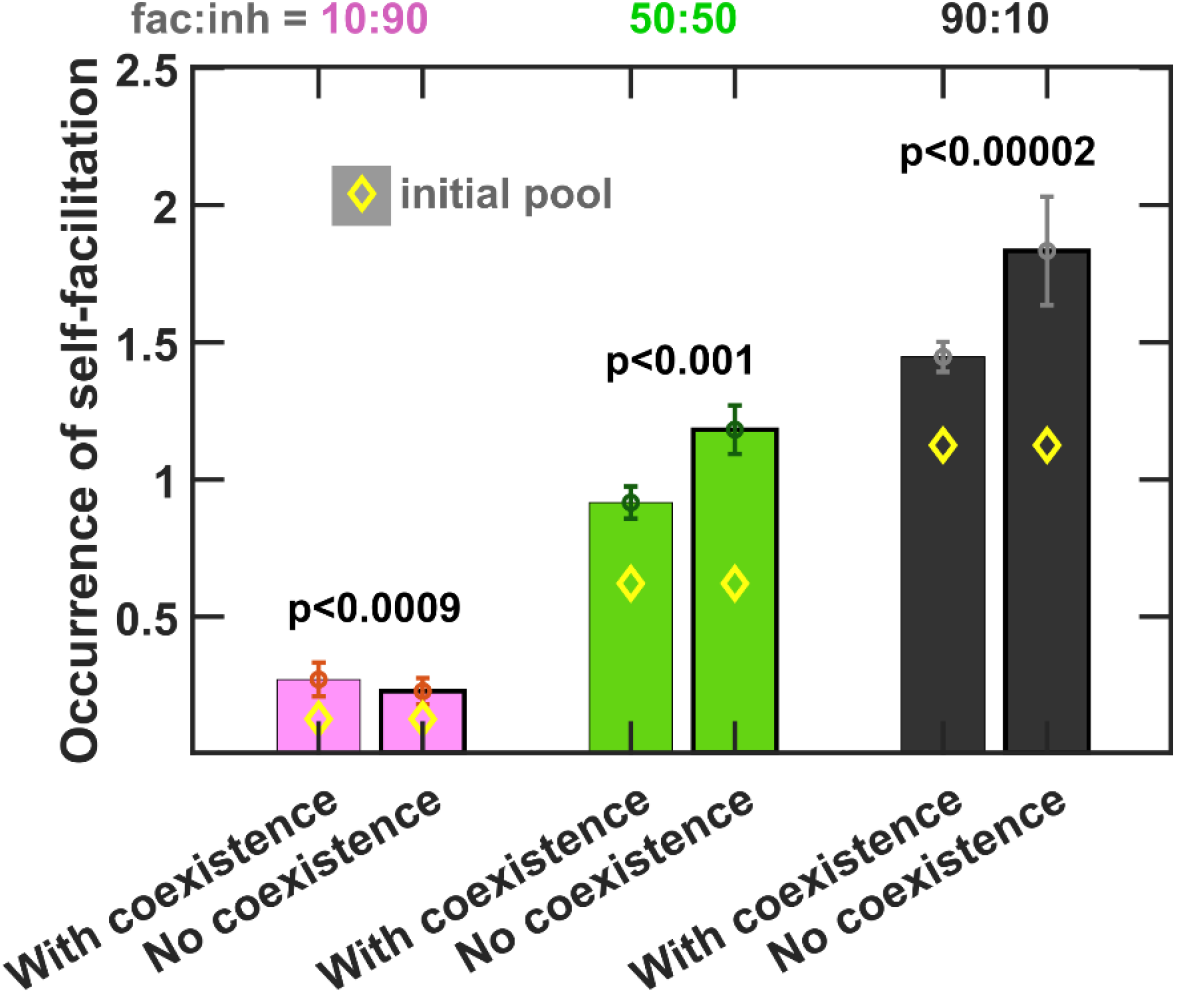
Self-facilitation is prominent among low-diversity outcomes. We separated the outcomes based on whether one (‘No coexistence’) or more species (‘With coexistence’) emerged from enrichment and examined the occurrence of self-facilitation. Occurrence is defined here as the average number of self-facilitation links (i.e. production of a mediator that benefits its producer) divided by the number of species present in the community. Each simulation started with 10 species and 5 mediators and ran for 100 generations. Here the average number of influences per mediator is 5 (*q*_*c*_ = 0.5) and the average number of mediators produced by each species is 2.5 (*q*_*p*_ = 0.5). Diamond markers (◊) show the occurrence in the initial pool of species before enrichment. The error bars are 95% confidence intervals generated by bootstrapping 100 samples using 500 total instances simulated. Mann-Whitney U test was used for comparison of occurrences between ‘No coexistence’ and ‘With coexistence’ cases to calculate the *p*-values. More self-facilitation occurrence was associated with absence of coexistence when fac:inh was 50:50 or 90:10, but the reverse trend held when interactions within the initial pool were mostly inhibitory (fac:inh = 10:90).

**Fig S8.**
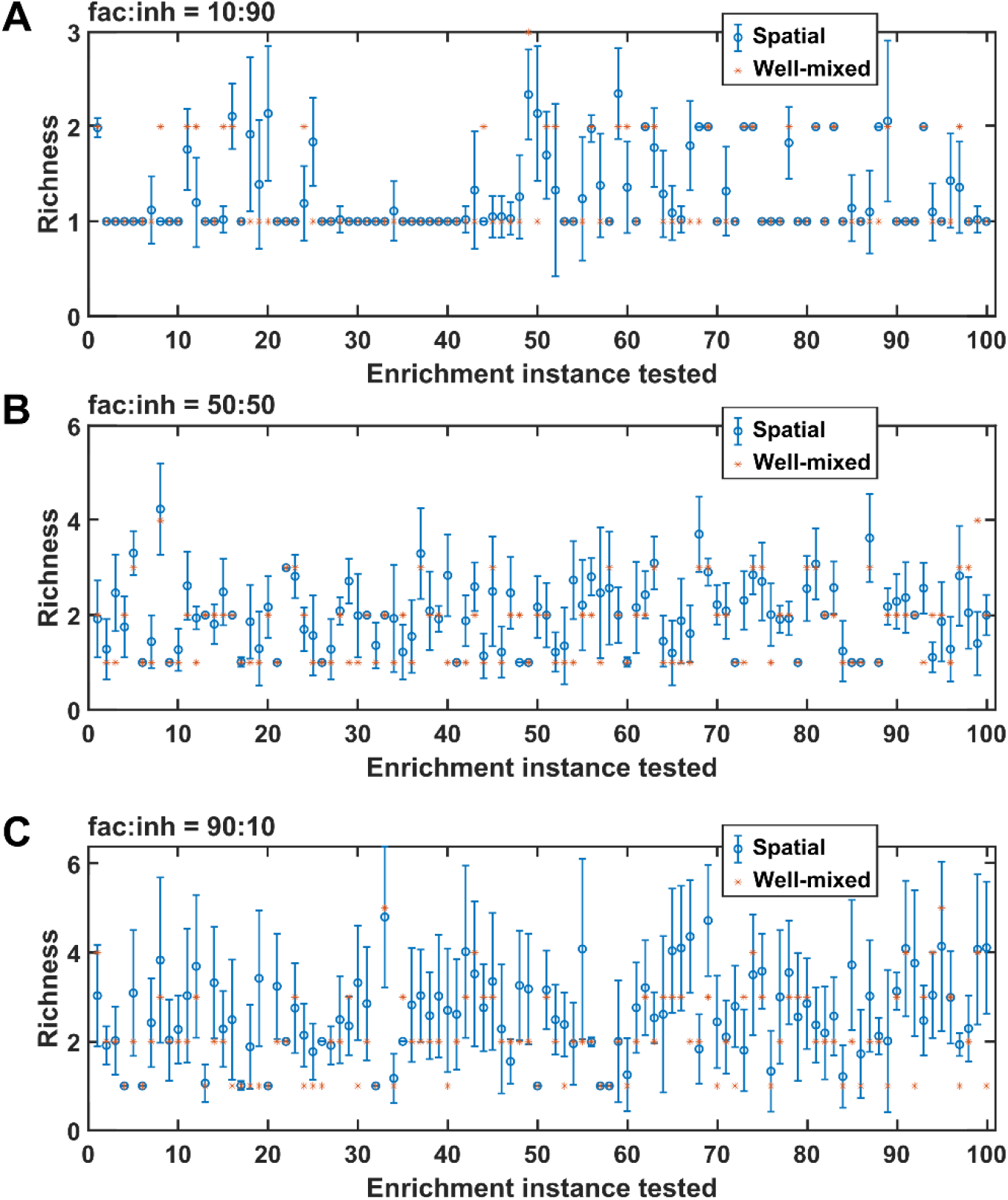
Rearranging the order of species affects spatial coexistence. For each of the cases of (A) fac:inh = 10:90, (B) fac:inh = 50:50, and (C) fac:inh = 90:10, we simulated 100 instances and in each instance, we examined coexistence outcome (final richness) in 100 permutations of the order of species (as an example, 10, 3, 6, 1, 5, 7, 4, 2, 9, 8 would be a permutation of the order of the 10 species). Asterisks (*) show the calculated richness for the corresponding well-mixed community, and the circle (o) and the error-bar show the mean and standard deviation of the richness values obtained for the 100 permutations tested. Overall, we observe that it is common to reach a different richness value when the order of species is changed.

**Fig S9.**
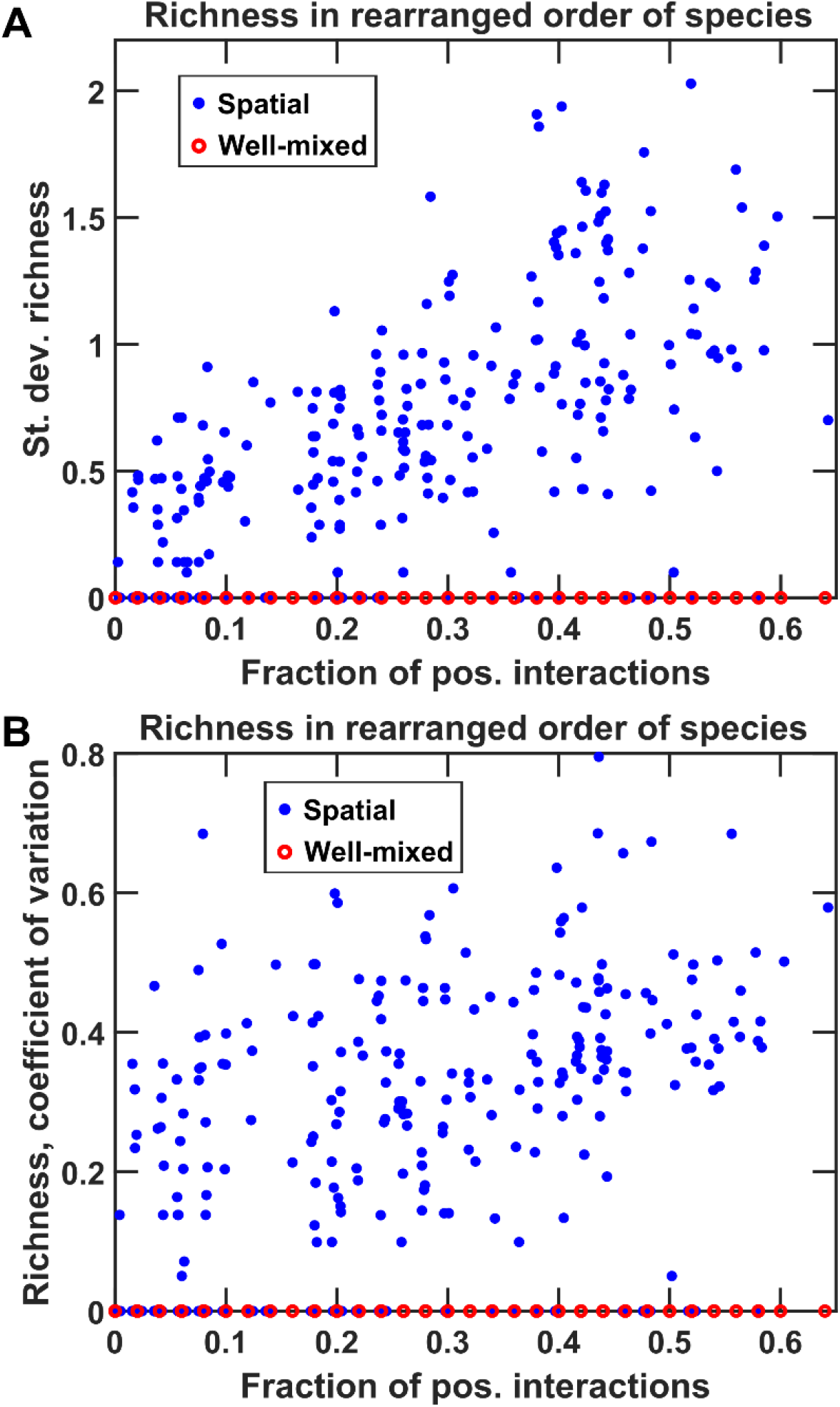
Rearranging the order of species affects spatial coexistence more in communities in which facilitation is prevalent. Similar to Fig S8, we simulated 100 instances each of fac:inh = 10:90, 50:50, and 90:10, and in each instance, we examined coexistence outcome (final richness) in 100 permutations of the order of species. Red circles (**○**) show the calculated richness for the corresponding well-mixed community, and the blue filled circles (●). The standard deviation of the richness values obtained for the 100 permutations are shown in (A). The trend was maintained for the coefficient of variations of the 100 permutations as well (B).

